# Cognate T-cell labeling by conventional and unconventional antigen-presenting cells *in vivo* using a modular SrtA-based toolkit

**DOI:** 10.64898/2026.09.17.752294

**Authors:** Guerric P.B. Samson, Yagmur Farsakoglu, Aurelie Lenaerts, Edit Horvath, Daniela Finke, Gleb Turchinovich

## Abstract

While different antigen-presenting cell (APC) populations can drive divergent T-cell fates even when presenting identical antigens (Ags), our ability to trace these specific *in vivo* interaction histories remains constrained by a lack of compatible molecular tools. Here, we describe an enhanced Labeling Immune Partnerships by SorTagging Intercellular Contacts (LIPSTIC) platform utilizing the immunologically inert human CD58/CD2 receptor-ligand pair (SrtA-hCD58/G5-hCD2). This system provides a modular alternative to previous LIPSTIC animal models and enables the detection and recovery of T cells following Ag-dependent interactions with distinct APC populations *in vitro* and *in vivo*. We applied this approach to interrogate the role of type 3 innate lymphoid cells (ILC3s), an unconventional RORγt⁺ Ag-presenting population whose role in shaping Ag-specific T cells *in vivo* remain poorly defined. Modified LIPSTIC successfully recorded cognate interactions between ILC3s and naive CD4⁺ T cells in the spleen and enabled direct comparison with CD4^+^ T cells activated by conventional dendritic cells (cDCs). Although interactions with ILC3s resulted in lower frequency of *in vivo*-labeled T cells than cDC2s, CD4^+^ T cells showed comparable cell division and expression of activation markers irrespectively of an APC type. Together, these findings establish SrtA-hCD58/G5-hCD2 as a versatile LIPSTIC receptor-ligand pair for tracking cell-cell interactions *in vivo* and provide a robust tool for studying how APC identity influences Ag-specific T-cell responses.

## Introduction

Cell-cell communication between antigen-presenting cells (APCs) and T cells is central to the regulation of adaptive immunity. Through antigen (Ag) presentation and the delivery of co-stimulatory and regulatory signals, APCs control T-cell activation, differentiation, and tolerance. Although conventional dendritic cells (cDCs) are considered the principal APCs within secondary lymphoid organs, accumulating evidence indicates that multiple APC populations contribute to the regulation of T-cell responses in distinct tissue environments. Among these populations, a family of RORγt⁺ APCs has emerged as an important regulator of adaptive immunity. In addition to RORγt⁺ extrathymic Aire-expressing cells (eTACs)^1–3^ and RORc⁺ DC-like cells^4,5^, type 3 innate lymphoid cells (ILC3s) can express MHC class II (MHCII) and directly interact with CD4⁺ T cells^6–8^. These findings suggest that Ag presentation and T-cell regulation extend beyond cDCs and may involve specialized APC populations with distinct immunological functions.

ILC3s are tissue-resident RORγt⁺ lymphocytes that contribute to mucosal immunity and tissue homeostasis^6–14^. Beyond these innate functions, ILC3s can process and present Ag via MHCII and directly regulate CD4⁺ T-cell responses. In the intestine, MHCII⁺ ILC3s maintain tolerance to commensal microbiota and dietary Ags by limiting microbiota-specific CD4⁺ T-cell responses^6,7^, whereas splenic MHCII⁺ ILC3s can prime Ag-specific CD4⁺ T cells and promote T-cell-dependent B-cell responses^15^. Together, these findings suggest that the outcome of ILC3/T-cell interactions is context dependent^16^, yet how Ag-specific interactions with ILC3s influence T-cell differentiation and fate remains poorly understood.

Studying APC/T-cell interactions and the fate of T cells following these interactions *in vivo* remains technically challenging. Although two-photon intravital microscopy enables visualization of immune cell contacts, it does not permit the isolation or molecular characterization of interacting cells following disengagement. Consequently, the molecular programs elicited by interactions with various APC populations remain difficult to define.

The recently developed Labeling Immune Partnerships by SorTagging Intercellular Contacts (LIPSTIC) approach addresses this challenge by using the bacterial enzyme sortase A (SrtA) to selectively label cells engaged in defined cell-cell interactions *in vivo*^17,18^. This strategy enables the subsequent identification, isolation, and molecular profiling of interacting cells by flow cytometry and single-cell RNA sequencing (scRNA-seq).

Here, we adapted and validated the LIPSTIC system using a new receptor-ligand pair and retroviral (RV)-based delivery. This approach demonstrated the Ag-specific labeling and recovery of CD4⁺ T cells following interactions with either cDC2s or ILC3s *in vivo.* Its modular design further enables adaptation to other immune cell types, providing a framework to investigate a broad range of cell-cell interactions *in vivo*.

## Methods

### Mice

C57BL/6J, B6.CD45.1/J (B6.SJL-Ptprc

<A> Pepc/BoyJ), C57BL/6JxB6.CD45.1/J (C57BL/6Jx B6.SJL-Ptprc

<A> Pepc/BoyJ), Rag2^-/-^Il-2rg^-/-^ (B6.129S6-Rag2<TM1FWA> x B6.129P2OlaHsd-Il2rg<TM1KRF>), CD45.1.SMARTA/J (B6.Tg(TcrLCMV)Aox; B6.SJL-Ptprc

<A>Pepc/BoyJ), CD45.1.SMARTA/J; Nur77-GFP (B6.Tg(TcrLCMV)Aox; Ptprc

<A> Pepc/BoyJ; Tg(Nr4a1-EGFP/cre)820Khog/J), CD45.1.OT-IIxRAG1 (C57BL/6-Ptprca-Tg(TcraTcrb)425Cbn x B6.129S7-Rag1tm1Mom/J), MHCII^-/-^ (B6.129S2-H2Ab1<TM1GRU>/J) and B6.OT2/Crl (C57BL/6-Tg(TcraTcrb)425Cbn/Crl) mice were bred and maintained under specific pathogen free (SPF) conditions in the animal facilities of the Department of Biomedicine (DBM; University of Basel, Switzerland). All animal experiments were conducted according to the Swiss Veterinary Law and Institutional Guidelines and were approved by the Cantonal Veterinary Office Basel City. Male or female mice aged 6-14 weeks were used for all experiments. Mice were housed under pathogen-free conditions in individually ventilated cages in a 22°C temperature-controlled room with 12 h light–12 h dark cycles and free access to food and water. Mice were euthanized by terminal CO_2_ inhalation.

### Cell lines and culture medium

Plat-E^19^ cells were cultured in Dulbecco’s Modified Eagle Medium (DMEM; 10566016, Thermo Fischer Scientific) supplemented with heat-inactivated 10% fetal calf serum (FCS; Gibco, Thermo Fischer Scientific), 1% penicillin/streptomycin (pen/strep; 15070063, Gibco), 50 µM β-mercaptoethanol (β-ME; 31350010, Gibco), 1x non-essential amino acids (NEAAs; 11140050, Gibco), and 1 mM sodium pyruvate (NaPyr; 11360070, Gibco). Interleukin (IL)-6 producing X63AG8.653 (X63) cells, granulocyte-macrophage colony-stimulating factor (GM-CSF) producing X63 cells, and stem cell factor (SCF) producing chinese hamster ovary (CHO) cells were maintained in IMDM complete medium consisting of IMDM (I3390-500ML, Sigma-Aldrich) supplemented with 2 mM l-glutamine (l–Glut; 25030081, Gibco), heat-inactivated 10% FCS, 5 mL insulin-transferrin-selenium-sodium pyruvate (51300044, Gibco), 1.5 mL 10% Primatone (P4963, Sigma-Aldrich), 1x NEAAs, 1% pen/strep, and 50 µM β-ME. Cells were seeded into a roller bottle and incubated for three to five days at 37°C. Supernatant was harvested, filtered, and stored at -20°C.

R10 medium consisted of RPMI 1640 medium (P04-18500, PAN-Biotech) supplemented with 10% heat-inactivated FCS, 1% pen/strep, and 50 µM β-ME.

### Generation of expression plasmids

Standard restriction cloning was performed using FastDigest restriction enzymes (Thermo Fisher Scientific) and the T4 DNA ligase (EL0011, Thermo Fisher Scientific). DNA assembly cloning was performed using the GenBuilder cloning kit (L00701, GenScript). PCR amplifications were performed using the Phusion Plus polymerase (F630S, Thermo Fisher Scientific). All plasmid DNA sequences were verified by Sanger sequencing (Microsynth).

The generation of the pMY-Puro-P2A-NUP98Hoxa10HD construct, was described elsewhere^20^.

For the pMY-flag-SrtA-CD40-IRES-GFP construct, an intermediate flag-SrtA-CD40 fragment was amplified by PCR from the pMP71-GFP-P2A-G5-myc-CD40 (Addgene plasmid #121166) and pMP71-Tomato-P2A-CD40L-SrtA-flag (Addgene plasmid #121167) plasmid, kindly provided by Gabriel Victora^17^. Next, the fragment was inserted into *XhoI* and *NotI* sites of a pMY-IRES-GFP plasmid.

For pMY-CD40L-myc-G5-IRES-mPlum construct, an intermediate CD40L-myc-G5 fragment was amplified by PCR from the pMP71-GFP-P2A-G5-myc-CD40 and pMP71-Tomato-P2A-CD40L-SrtA-flag. Next, the fragment was inserted into *EcoR1* and *NotI* sites of a pMY-IRES-mPlum plasmid.

For the pMY-flag-SrtA-hCD58-IRES-GFP construct, an intermediate flag-SrtA-hCD58 fragment was amplified by PCR from the pMY-flag-SrtA-CD40-IRES-GFP and the CD58 (NM_001144822) Human Tagged ORF Clone Lentiviral Particle (RC227634, Origene). Next, the fragment was inserted into *XhoI* and *NotI* sites of a pMY-IRES-GFP plasmid.

For pMY-G5-myc-hCD2-IRES-mPlum construct, an intermediate G5-myc-hCD2 fragment was amplified by PCR from the pMY-CD40L-myc-G5-IRES-mPlum and the pcDNA3-hCD2 plasmid, kindly gifted from David Ron (Addgene plasmid # 21810). Next, the fragment was inserted into *EcoR1* and *NotI* sites of a pMY-IRES-mPlum plasmid.

For the pMP71-GFP-P2A-flag-SrtA-hCD58 construct, the flag-SrtA-hCD58 fragment was amplified by PCR from the pMY-flag-SrtA-hCD58-IRES-GFP plasmid and subsequently assembled with the pMP71-GFP-P2A-G5-myc-CD40 plasmid using a DNA assembly reaction.

For the pMP71-Tomato-P2A-G5-myc-hCD2 construct, the G5-myc-hCD2 fragment was amplified by PCR from the pMY-G5-myc-hCD2-IRES-mPlum plasmid and subsequently assembled with the pMP71-Tomato-P2A-CD40L-SrtA-flag plasmid using a DNA assembly reaction.

Plasmids and plasmid maps are available upon request.

### Production of recombinant viral particles

Retroviral particles were generated by transfecting Plat-E cells using Polyethylenimine (PEI; 23966, Polyscience) with retroviral expression plasmids pMY-Puro-P2A-NUP98Hoxa10HD, pMP71-GFP-P2A-G5-myc-CD40, pMP71-Tomato-P2A-CD40L-SrtA-flag, pMP71-GFP-P2A-flag-SrtA-hCD58, pMP71-Tomato-P2A-G5-myc-hCD2, pMY-G5-myc-hCD2-IRES-mPlum, pMY-flag-SrtA-hCD58-IRES-GFP, pMY-flag-SrtA-CD40-IRES-GFP, pMY-CD40L-myc-G5-IRES-mPlum. Supernatant containing retroviral particles was collected at 48 and 72 hours (h) post-transfection and filtered (0.45 μm, Polyethersulfone (PES) membrane; 83.1826, Sarstedt).

### Primary cell isolation and culture

Bone marrow (BM) was flushed from the femur and tibia of 6-9-week-old B6.CD45.1/J or CD45.1.SMARTA/J mice. Cells expressing lineage markers were then depleted by first labeling them with a mixture of PE-conjugated antibodies (Abs) targeting lineage-specific markers (CD3, CD4, CD8, CD19, B220, Gr-1, TER119, F4/80, CD11b and CD11c), followed by anti-PE microbead-based depletion (130-048-801, Miltenyi Biotech). Lineage negative (lin^−^) BM cells were cultured in IMDM complete medium containing saturating amounts of SCF and IL-6 supernatant for two days prior immortalization with NUP98Hoxa10HD (NUPA10hd).

To isolate splenocytes or splenic DCs, spleens were *ex vivo* injected with 500 μL digestion solution [R10 medium supplemented with 400 ng/mL collagenase D and 0.004 mg/mL DNase I], incised using scissors, and subsequently incubated in 2 mL digestion solution for 30 min at 37°C. The resulting cell suspension was applied to a 70 µm cell strainer. If needed, DCs were enriched using CD11c microbeads for mouse (130-125-835, Miltenyi Biotec) and sorted by flow cytometry. Some purified cDC2s (CD11c^+^MHCII^+^CD11b^+^) were then matured overnight in R10 medium supplemented with 100 ng/mL LPS, 200 ng/mL Flt3L (300-19, PeproTech, Thermo Fisher Scientific) and 20 ng/mL GM-CSF.

To isolate small intestine (SI) cells from lamina propria, the SI was opened longitudinally, cut into pieces and washed twice with 10 mM HEPES, 5 mM Ethylenediaminetetraacetic acid (EDTA), 5% FCS in 1x PBS for 20 min at 37°C and once with 1x PBS for 20 min to remove feces and mucus. Tissue pieces were digested in DMEM supplemented with 10 mM HEPES, 5% FCS, 0.025 mg/mL DNase I (10104159001, Roche), 1 mg/mL collagenase D (11088858001, Sigma Aldrich) and 0.5 mg/mL Dispase (D4693, Sigma Aldrich) in gentleMACS™ C tubes (130-093-237, Miltenyi Biotec) at 37°C for 35 min (rotating). Next, tissue pieces were dissociated by running gentleMACS program m_intestine_01 on a gentleMACS™ Dissociator (Miltenyi Biotec) to obtain a cell suspension. Supernatant was collected through a 70 µm strainer. SI cells were purified by Percoll density gradient (40% /80%) centrifugation.

SI- or splenic-derived ILC3s were sorted from the lamina propria or from the spleen, respectively, of RAG1^-/-^ mice as Thy1.2^+^NK1.1^-^KLRG1^-^ Lin^-^ (lin cocktail consisted of Abs against CD3, CD8, TCRβ, TCRγδ, CD19, B220, Ter119, CD11b, CD11c, F4/80 and Gr-1) and cultured in complete RPMI 1640 medium supplemented with 2 mM l-Glut, 10% heat-inactivated FCS, 1% pen/strep, 1x NEAAs, 1 mM NaPyr, 10 mM HEPES (15630056, Gibco) and 50 µM β-ME in the presence of 20 ng/mL IL-2 (212-12, PeproTech), 20 ng/mL IL-7 (217-17, PeproTech), 20 ng/mL SCF (250-03, PeproTech) and 1 µM *all*-*trans*-retinoic acid (RA; R2625, Sigma Aldrich) for 14 days. During the final four days of culture, medium was supplemented with 40 ng/mL IFNγ (575306, BioLegend) to induce MHCII expression. Cells were then sorted for MHCII⁺ ILC3s and cultured overnight in the presence of 20 ng/mL IL-1β (575102, BioLegend) to induce maturation.

To isolate naive CD4^+^ T cells, the spleen was smashed through a 70 µm cell strainer, washed with PBS, and splenocytes were resuspended in PBE buffer (PBS + 2% FCS + 2 mM EDTA). Subsequently, naive CD4^+^ T cells were isolated using the EasySep™ Mouse naive CD4^+^ T-cell isolation kit (19765, STEMCELL) according to the manufacturer’s protocol. T cells were maintained in RPMI 1640 medium supplemented with 2 mM l-Glut, 10% heat-inactivated FCS, 1% pen/strep, 1x NEAAs, 1 mM NaPyr, 10 mM HEPES and 50 µM β-ME.

Lymph nodes (LNs) were harvested and incubated in 1 mL digestion solution [2 mg/mL collagenase D and 0.004 mg/mL DNAse I in R10] for 45 min at 37°C, smashed through a 70 µm cell strainer and washed with PBE buffer. BM was isolated by crushing the bones in a mortar and pestle. Cells were filtered through a 70 µm cell strainer and washed with PBS, erythrocytes were lysed, and cells were resuspended in R10 medium.

Thymi were smashed through a 70 µm cell strainer and washed with PBE buffer.

### Flow cytometry

Single-cell suspensions derived from mouse tissue were incubated with anti-FcγRII/RIII Ab supernatant of clone 2.4G2 and surface marker-specific Abs for 40 min at 4°C in PBE Buffer. Live/dead staining was conducted using the fixable viability dye Zombie Aqua™ (423101, BioLegend). For the intracellular staining of transcription factors, cells were fixed and permeabilized with the Foxp3 transcription factor staining buffer set (00-5523-00, Thermo Fisher Scientific) according to the manufacturer’s instructions. Data were acquired on an LSRFortessa (Waters Biosciences), CytoFLEX (Beckman Coulter), or Aurora (Cytek). Cell sorting was conducted using a FACS Aria II (Waters Biosciences). The Diva software (BD FACS Aria II and BD LSRFortessa), CytExpert software (CytoFLEX, Beckman Coulter), and SpectroFlow software (Aurora, Cytek) were used for data collection. Data were analyzed using the FlowJo™ v10.9 software (Waters Biosciences). The full list of Abs is available in **Table 1**.

### Generation and maintenance of NUPA10hd progenitor cells

The transduction of lin^−^ BM cells with pMY-Puro-P2A-NUP98Hoxa10HD containing viral particles as well as the genetic engineering of NUPA10hd immortalized progenitor cells with LIPSTIC constructs has been previously described^20^. Cells were maintained in IMDM complete medium containing saturating amounts of SCF and IL-6. NUPA10hd progenitors were maintained in culture for at least 12 weeks.

### Generation of DCs from NUPA10hd progenitor cells

B6.CD45.1/J NUPA10hd progenitors were washed three times with R10 medium to remove SCF and IL-6 and 50’000 cells (per non-treated 10 cm petri dish; 633180, Greiner) were differentiated into immature DCs (iDCs) in R10 medium supplemented with 20 ng/mL GM-CSF from conditioned supernatant (determined by ELISA) for 9 days. iDCs were matured with 100 ng/mL Lipopolysaccharide (LPS) from Escherichia coli O111:B4 (L4391, Sigma-Aldrich) for 24 h. Prior to functional assays, mature DCs (mDCs) were enriched for CD11c using CD11c microbeads for mouse (130-125-835, Miltenyi Biotec).

### Reconstitution of RAGγc^-/-^ or CD45.1/CD45.2 mice with NUPA10hd progenitors

CD45.2^+^ Rag2^-/-^ Il2rg^-/-^ (RAGγc^-/-^) mice were sub-lethally irradiated with 450 cGy, whereas C57BL/6JxB6.CD45.1/J (CD45.1/CD45.2) mice received two consecutive 500 cGy doses administered 4 h apart. The next day, 1x10^7^ NUPA10hd progenitors were injected intravenously (*i.v.)* into RAGγc^-/-^ or CD45.1/CD45.2 mice. Mice were sacrificed at 4 to 6 weeks after cell transfer.

### LIPSTIC retroviral transduction and cell-cell interaction

For 291PC cells, stable transduction with the indicated expression vectors was performed using the same protocol as described for NUPA10hd progenitors^20^. 291PCs expressing G5- or SrtA-fusion constructs were mixed at 1:1 ratio (0.5x10^6^ cells of each population) in a 1.5 mL conical tube, to which biotin-LPETG was added to a final concentration of 100 μM. Cells were incubated at room temperature for 30 min and washed three times with PBE to remove excess biotin-LPETG before FACS staining.

Splenic- and SI-derived ILC3s were transduced with retroviral particles containing GFP and SrtA-hCD58, following a protocol previously described^21^, and sorted for GFP^+^ and Thy1.2^+^ cells four days later. NUPA10hd mDCs expressing G5- or SrtA-fusion constructs were generated from progenitor cells *in vitro*.

Naive CD4^+^ T cells isolated from B6.OT2 mice were activated with anti-CD3/anti-CD28 Dynabeads^TM^ (11452D, Thermo Fisher Scientific) for 24 h and transduced with retroviral particles containing CD40L-SrtA or G5-hCD2, as previously described^21^. Three days later, transduced CD4^+^ T cells were sorted for Tomato^+^ cells and maintained in a resting state in T-cell medium supplemented with 10 ng/mL IL-7 and 20 IU/mL IL-2. To generate naive G5-hCD2^+^ SMARTA CD4^+^ T cells, lethally irradiated CD45.1/CD45.2 mice were reconstituted with G5-hCD2^+^ NUPA10hd SMARTA progenitors. 6 weeks later, naive G5-hCD2^+^ CD4^+^ T cells were isolated from the spleen. Mature SrtA^+^ ILC3s and NUPA10hd mDCs expressing G5- or SrtA-fusion constructs were loaded with 1 μM OVA_323–339_ (cognate; O1641, Sigma-Aldrich) or LCMV-GP_61–80_ (control; GeneCust) peptide for 2 h at 37 °C. Cells were then washed three times in R10 medium and seeded into a U-bottom 96-well plate together with rested SrtA^+^ or G5^+^ OT-II CD4^+^ T cells in a 1:1 ratio (total of 60’000 cells) for 6 h. Biotin-LPETG was added in the last 20 min of co-culture at a final concentration of 10 µM. Some CD4^+^ T cells were incubated with an αCD40L blocking Ab (BE0017-1, Bio X Cell) or an isotype Ab (BE0091, Bio X Cell) for 30 min before the beginning of co-culture at a final concentration of 150 μg/mL. At the end of co-culture, cells were washed three times with PBE before FACS staining to remove excess biotin-LPETG substrate.

For *in vivo* LIPSTIC in popliteal LNs (PLNs), SrtA-hCD58^+^ splenic cDCs were loaded with 5 μM OVA_323–339_ (cognate) or LCMV-GP_61–80_ (control) peptide. 1x10^6^ cDCs (10 μL) were injected subcutaneously (*s.c.*) together with 0.4 μg/mL LPS into the hind hock of MHCII^-/-^ mice. 24 h later, 1x10^6^ rested G5-hCD2^+^ OT-II CD4^+^ T cells were *i.v*. injected. 10 h later, mice were injected three times at 45 min intervals with biotin-LPETG (10 μL of 20 mM solution in PBS) *s.c.* into the hind hock, and PLNs were collected 40 min after the last injection.

For *in vivo* LIPSTIC in spleen, 1x10^6^ mature SrtA-hCD58^+^ ILC3s or splenic cDC2s loaded with 5 μM OVA_323–339_ (control) or LCMV-GP_61–80_ (cognate) peptide were injected *i.v.* into MHCII^-/-^ mice. 24 h later, 0.5x10^6^ freshly sorted naive G5-hCD2^+^ SMARTA CD4^+^ T cells (CD4^+^CD62L^+^CD44^-^Tomato^hi^) were injected *i.v*. 10 h later, mice were injected intraperitoneally (*i.p)* three times at 45 min intervals with biotin-LPETG (100 μL of 50 mM solution in PBS). Spleen and inguinal LNs (iLNs) were collected 40 min after the last injection.

Biotin-aminohexanoic acid**-**LPETGS (C-term amide, 95% purity) was purchased from LifeTein and stock solutions were prepared in PBS at 20 or 50 mM (stored at -80°C).

### T-cell proliferation assay *in vivo*

To test T-cell proliferation *in vivo*, 1x10^6^ mature cDC2s or ILC3s were loaded with 5 μM LCMV-GP_61–80_ peptide and injected *i.v.* into MHCII^-/-^ recipient mice. The next day, naive CD45.1^+^ Nur77-GFP SMARTA CD4^+^ T cells were labeled with CellTrace Violet (CTV; C34557, Thermo Fischer Scientific) following the manufacturer’s guidelines, and 0.5x10^6^ cells were injected *i.v*. Two days later, mice were sacrificed and spleens were harvested. The proliferation index was calculated based on CTV dilution profiles by dividing the total number of cell divisions by the number of cells that went into division.

### Confocal microscopy

1x10^6^ mature SrtA-hCD58^+^ ILC3s were injected *i.v.* into MHCII^-/-^ mice. Two days later, mice were sacrificed, and the right ventricle was opened to allow outflow during perfusion. The mice were then perfused through the left ventricle with 10 mL PBS, followed by 10 mL 1% PFA (28908, Thermo Fisher Scientific), prior to spleen collection. Spleens were cut in half and fixed in 1% PFA for 24 h, and embedded in 6% low-gelling agarose (A9045, Sigma Aldrich). Spleens were cut on a vibratome (80-140 μm) and sections were blocked with CD16/CD32 (clone 93; 101302, BioLegend) in staining buffer (PBS + 1% FCS + 0.1% Tween 20 + 0.05% NaN_3_) for three days at 4°C. Sections were then stained for one week at 4°C with Abs to identify B cell follicles (IgD; clone 11-26c.2a; 405708, BioLegend), marginal zone (CD169; clone 3D6.112; 142421, BioLegend), red pulp (F4/80; clone BM8; 123110, BioLegend) and GFP (A-11122, Thermo Fisher Scientific) in staining buffer, washed with staining buffer and stained with a goat anti-rabbit IgG (H+L) Ab, Alexa Fluor 488-conjugated (A-11008, Thermo Fisher Scientific) in staining buffer for three days at 4°C. After washing with staining buffer, spleen sections were mounted with Fluoromount^TM^ aqueous mounting medium (F4680, Sigma Aldrich) and images were acquired on a Leica Stellaris 8 Falcon confocal microscope using a 20x air objective (z-stacks of 3-µm step size). Imaris (Bitplane) was used for analysis.

## Results

### Engineering of an immunologically inert receptor-ligand pair for LIPSTIC labeling

A previous study has shown that the LIPSTIC G5-CD40/CD40L-SrtA receptor-ligand pair enables the labeling of G5-CD40^+^ DCs (acceptors) that have engaged in transient interactions with CD40L-SrtA^+^ T cells (donors) via the transfer of biotin-labeled LPETG peptide (LIPSTIC labeling) from donor to acceptor cells^17^. However, to determine how APC identity affects the fate of an interacted T-cell *in vivo*, an inverse approach is required where APCs act as donors and T cells act as acceptors. We first generated and validated the inverted LIPSTIC strategy, with SrtA-CD40 as the donor and CD40L-G5 as the acceptor in 291PC cells (**Figure 1A**). Biotin-labeled LPETG was efficiently transferred from donor to acceptor cells, confirming the functionality of the inverted configuration (**Figure 1B**). Notably, CD40L-G5^+^ 291PC cells labeling was inhibited by an αCD40L blocking antibody, confirming that LIPSTIC labeling was dependent on the specificity of the engineered receptor-ligand interaction (**Figure 1C**). Because constitutive overexpression of costimulatory molecules may lead to broad immune activation, while physiological regulation of expression would require the generation of knock-in mice, we established an “universal”, immunologically inert LIPSTIC receptor-ligand pair. We selected the human CD58/CD2 receptor-ligand pair because it does not cross-interact with its murine homologues. We generated and validated the SrtA-hCD58/G5-hCD2 receptor-ligand pair, with efficient LIPSTIC labeling of acceptor cells by donor cells in 291PC co-cultures (**Figure 1D-E**). Importantly, we observed a similar mean fluorescence intensity (MFI) for the biotin-LPETG signal of acceptor G5-hCD2^+^ and CD40L-G5^+^ 291PC cells, indicating a similar labeling efficiency by their respective donors (**Figure 1F**). Furthermore, when co-culturing acceptor 291PC cells with mismatched receptor donor pairs, we observed no increase in the MFI of biotin-LPETG on the acceptor cells, indicating that each receptor could only transfer the biotin-LPETG substrate specifically to its respective ligand (**Figure 1G-H**).

**Figure 1:**
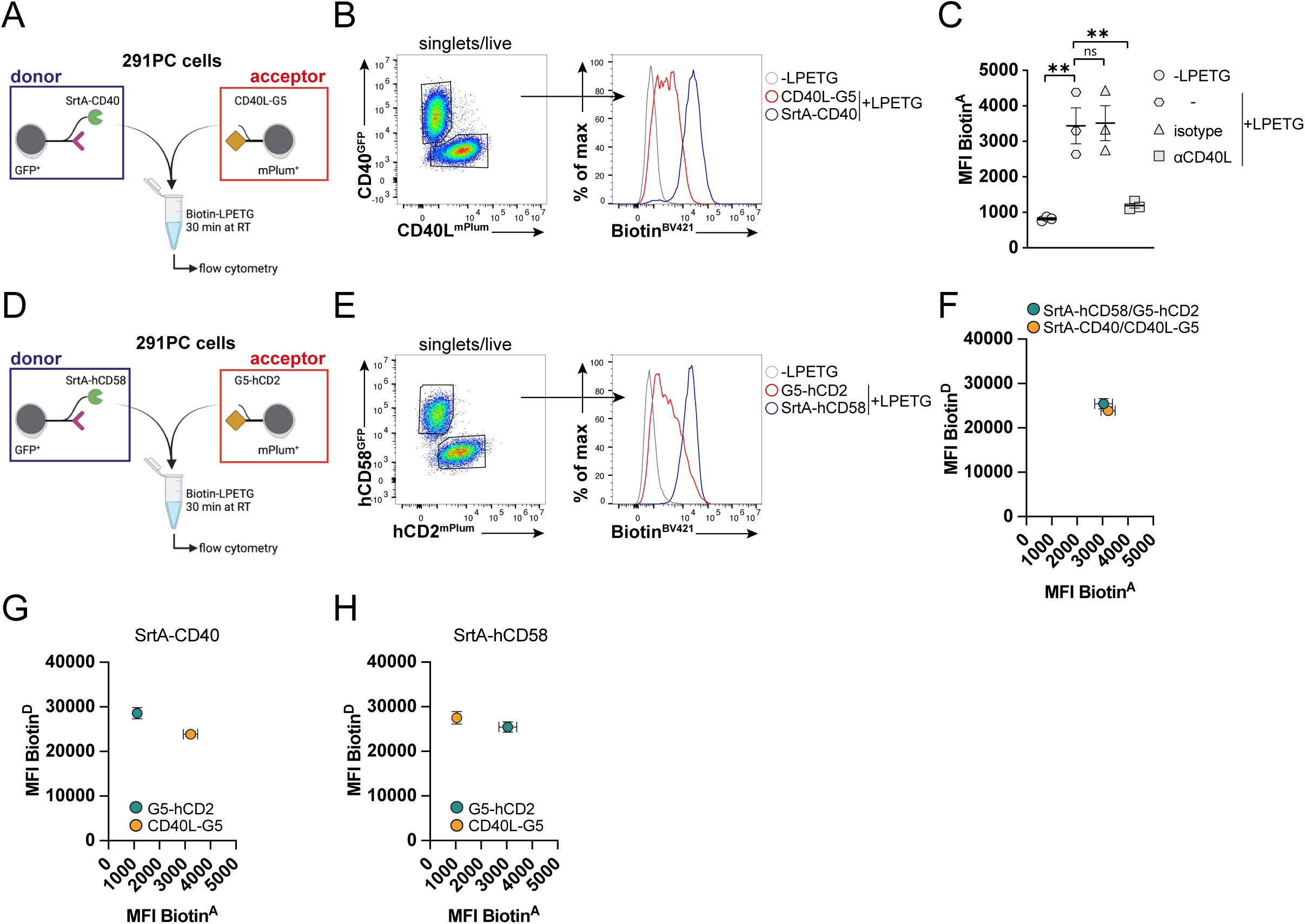
Generation and validation of a LIPSTIC hCD58/hCD2 receptor-ligand pair. Scheme of the LIPSTIC approach (**A**) and flow cytometry analysis (**B**) showing biotin staining in SrtA-CD40^+^ (D; donor) and CD40L-G5^+^ (A; acceptor) 291PC cells. (**C**) Intercellular labeling of CD40L-G5^+^ 291PC cells pre-treated or not with a blocking αCD40L or isotype antibody. Scheme of the LIPSTIC approach (**D**) and flow cytometry analysis (**E**) showing biotin staining in SrtA-hCD58^+^ (D) and G5-hCD2^+^ (A) 291PC cells. (**F**) Quantification of the LIPSTIC labeling using SrtA-hCD58/G5-hCD2 and SrtA-CD40/CD40L-G5 expressing 291PC cells. Quantification of the LIPSTIC labeling using SrtA-CD40 (**G**) or SrtA-hCD58 (**H**) and G5-hCD2 or CD40L-G5 expressing 291PCs. Mean values ± SEM of three independent experiments. Statistical significance was determined by one-way ANOVA, followed by Tukeys multiple-comparisons test. p>0.05 (ns; not significant) and p<0.01 (**).

### hCD58/hCD2 LIPSTIC enables monitoring of Ag-specific interactions between DCs and T cells *in vitro* and *in vivo*

To measure LIPSTIC labeling during Ag-specific interactions between APCs and T cells , we established an efficient system to immortalize and genetically modify hematopoietic BM progenitors using the fusion protein NUP98Hoxa10HD (NUPA10hd)^22–25^. We recently showed that wild-type (WT) and genetically engineered NUPA10hd progenitors efficiently generated DCs *in vitro* and retained the ability to reconstitute myeloid and lymphoid lineages *in vivo*^20^. As proof of concept for LIPSTIC labeling, we generated SrtA-hCD58^+^ mature DCs (mDCs) *in vitro* from NUPA10hd progenitors of WT and G5-hCD2^+^ CD4^+^ T cells from TCR transgenic OT-II mice (**Figure 2A**). Following a protocol summarized in **Figure 2B**, we confirmed biotin-LPETG loading on donor mDCs (**Figure 2C-D**) and significantly increased intercellular labeling when G5-hCD2^+^ CD4^+^ T cells (acceptors) were co-cultured with OVA_323-339_-loaded SrtA-hCD58^+^ mDCs (donors) compared to the control LCMV-GP_61-80_-treated group (**Figure 2C,E**). Notably, we observed an increase in CD69 expression within biotin^+^ G5-hCD2^+^ OT-II CD4^+^ T cells cultured with OVA_323-339_-loaded SrtA-hCD58^+^ mDCs, compared to the control LCMV-GP_61-80_-treated group (**Figure 2C,F**), facilitating the identification of Ag-specific interacting T cells. Intercellular labeling was also observed with the original CD40/CD40L LIPSTIC pair^17^ when CD40L-SrtA^+^ CD4^+^ T cells (donors) were co-cultured with OVA_323–339_-loaded G5-CD40^+^ mDCs (acceptors) (**Figure S1A-G**).

**Figure 2:**
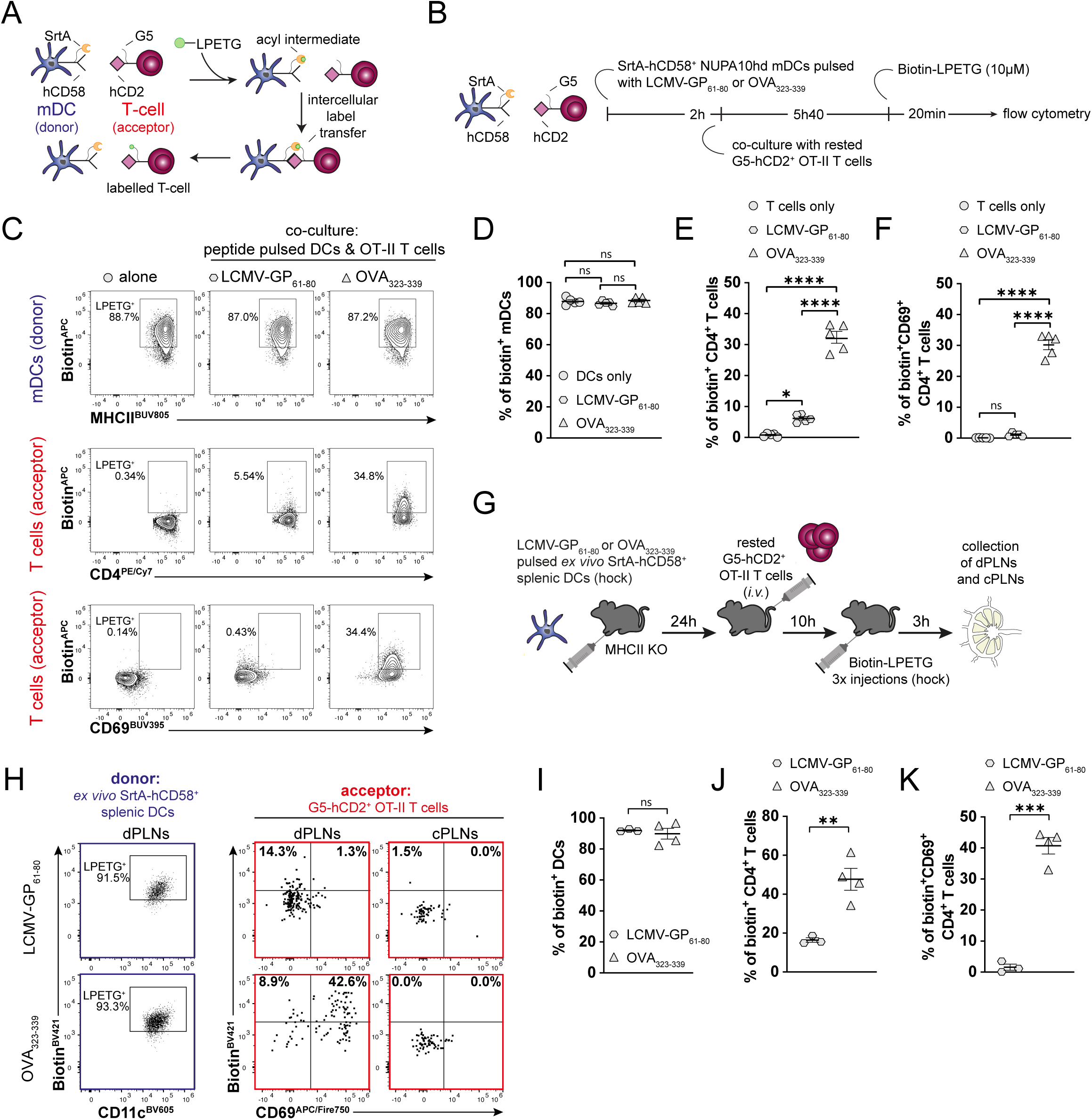
Labeling of SrtA-hCD58^+^ DCs-primed G5-hCD2^+^ T cells *in vitro* and *in vivo*. (**A**) Scheme of the modified LIPSTIC approach. (**B**) Experimental setup to analyze intercellular labeling of G5-hCD2^+^ OT-II CD4^+^ T cells when co-cultured with 1 µM OVA_323-339_ or LCMV-GP_61-80_-loaded SrtA-hCD58^+^ mature DCs (mDCs) by FACS *in vitro*. (**C**) Representative flow cytometry contour plots depicting the loading of biotin-LPETG on SrtA-hCD58^+^ mDCs (top) and its transfer to G5-hCD2^+^ T cells (middle-bottom). Frequencies of biotin^+^ mDCs (**D**), biotin^+^ (**E**) and biotin^+^CD69^+^ (**F**) CD4^+^ T cells. Mean values ± SEM of five independent experiments. Statistical significance was determined by one-way ANOVA, followed by Tukeys multiple-comparisons test. p>0.05 (ns; not significant), p<0.05 (*) and p <0.0001 (****). (**G**) Scheme of the experimental setup for the LIPSTIC labeling of G5-hCD2^+^ T cells by SrtA-hCD58^+^ *ex vivo* splenic DCs in draining popliteal lymph nodes (dPLNs). (**H**) Representative flow cytometry dot plots depicting the loading of biotin-LPETG on SrtA-hCD58^+^ DCs (left) and its transfer to G5-hCD2^+^ T cells in the draining (middle) or contralateral (right) PLNs. Frequencies of biotin^+^ DCs (**I**), biotin^+^ (**J**) and biotin^+^CD69^+^ (**K**) CD4^+^ T cells in the dPLNs. Mean values ± SEM of 3-4 mice from two independent experiments. Statistical significance was determined by unpaired Welch’s Test. p>0.05 (ns; not significant), p<0.01 (**) and p <0.001 (***).

To determine whether our LIPSTIC approach could be applied *in vivo*, we first generated SrtA-hCD58⁺ splenic DCs by reconstituting RAGγc^-/-^ mice with retrovirally transduced SrtA-hCD58⁺ NUPA10hd progenitors (**Figure S2A**). Irradiated RAGγc^-/-^ mice reconstituted with these progenitors displayed normal frequencies of lymphoid and DC progenitor populations, spleen size, and functional splenic GFP^+^ biotin^+^ DCs upon *ex vivo* incubation with the biotin-LPETG substrate (**Figure S2B-F**). Next, OVA_323-339_-loaded *ex vivo* SrtA-hCD58^+^ DCs were injected *s.c.* into the hind hock of MHCII^-/-^ recipient mice, and G5-hCD2^+^ OT-II CD4^+^ T cells were injected *i.v.* 24 hours (h) later. The biotin-LPETG substrate was administered by three injections between 10 h and 13 h after T-cell transfer (**Figure 2G**). Flow cytometry of draining popliteal lymph node (dPLN) cells showed efficient and Ag-peptide-specific LIPSTIC labeling of transferred OT-II T cells by OVA_323-339_-loaded splenic DCs (**Figure 2H-K**). As expected, no LIPSTIC labeling of T cells occurred in the contralateral PLNs (cPLNs) (**Figure 2H; right**). Taken together, we successfully generated a novel versatile LIPSTIC receptor-ligand pair that enabled the tracking of CD4⁺ T cells following cognate interactions with DCs *in vitro* and *in vivo*.

### *Ex vivo-*expanded SrtA^+^ ILC3s present Ag and efficiently label T cells *in vitro*

Although cDCs are well-characterized professional APCs, accumulating evidence points to a broader family of unconventional RORγt⁺ APCs, such as ILC3s, that can regulate CD4⁺ T-cell responses^4–7,15,16^. To investigate how T-cell interactions with unconventional APCs differ from those with conventional APCs, we applied our human CD58/CD2 LIPSTIC system to ILC3s. We established an *ex vivo* culture protocol for ILC3s using stem cell factor (SCF), interleukin (IL)-2, IL-7, and *all*-*trans*-retinoic acid (RA), which facilitated efficient expansion and subsequent retroviral transduction of splenic- and small intestine (SI)-derived ILC3s with SrtA-hCD58 (**Figure 3A**).

**Figure 3:**
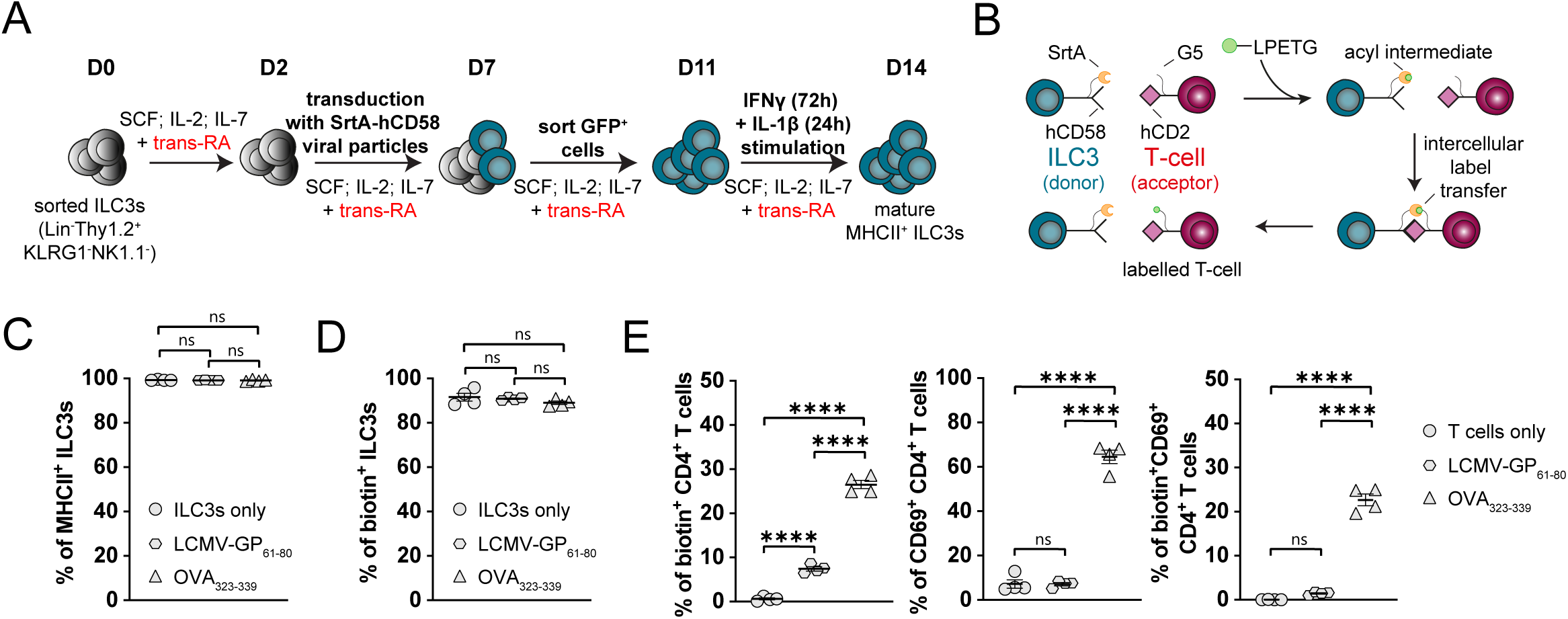
*Ex vivo*-expanded Ag-presenting SrtA^+^ ILC3s specifically label and activate cognate CD4^+^ T cells. (**A**) Scheme of the *ex vivo* ILC3 expansion protocol. (**B**) Experimental setup to analyze intercellular labeling of G5-hCD2^+^ OT-II CD4^+^ T cells when co-cultured with 1 µM OVA_323-339_ or LCMV-GP_61-80_-loaded mature SrtA-hCD58^+^ ILC3s by FACS *in vitro*. Frequencies of MHCII^+^ (**C**) and biotin^+^ (**D**) mature ILC3s. (**E**) Frequencies of biotin^+^ (left), CD69^+^ (middle) and biotin^+^CD69^+^ (right) CD4^+^ T cells. Mean values ± SEM of four independent experiments. Statistical significance was determined by one-way ANOVA, followed by Tukeys multiple-comparisons test. p>0.05 (ns; not significant) and p <0.0001 (****).

Similar to DCs, ILC3s acquire full Ag-presenting capacity through a maturation process involving the upregulation of MHCII and co-stimulatory molecules in response to interferon gamma (IFNγ) and IL-1β stimulation, respectively^15,16^ (**Figure 3A**). Here, we defined ILC3s cultured with IFNγ and IL-1β as mature. Interestingly, no major differences in MHCII or different co-stimulatory molecule expression were observed between *ex vivo* cultured splenic- and SI-derived ILC3s upon maturation (**Figure S3**). Given their greater abundance in the SI, we used *ex vivo*-expanded and matured SI-derived ILC3s for subsequent experiments. To measure LIPSTIC labeling during Ag-specific ILC3/T-cell interactions *in vitro*, mature SrtA-hCD58^+^ ILC3s were loaded with either the cognate OVA_323–339_ or with a control LCMV-GP_61–80_ peptide and co-cultured with G5-hCD2^+^ OT-II CD4^+^ T cells (**Figure 3B-C**). Biotin-LPETG efficiently bound to SrtA on MHCII^+^ ILC3s (**Figure 3D**) and was subsequently transferred onto G5-hCD2^+^ CD4^+^ T cells in an Ag-specific manner (**Figure 3E; left-middle**). Notably, the extent of LIPSTIC labeling and T-cell activation, measured by biotin and CD69 expression (**Figure 3E; right**), was comparable to that observed with DCs (**Figure 2F**). Taken together, we established a robust ILC3 *ex vivo* expansion protocol that allows us to generate Ag-presenting ILC3s that efficiently label cognate CD4⁺ T cells *in vitro*.

### *Ex vivo*-expanded mature ILC3s localize to the splenic white pulp-marginal zone border and induce CD4^+^ T-cell proliferation

To assess whether *ex vivo-*expanded mature ILC3s remain functional *in vivo*, we examined their ability to home to the spleen and induce CD4^+^ T-cell activation and proliferation following adoptive transfer. First, we transferred mature SrtA-hCD58^+^ ILC3s *i.v.* into MHCII^-/-^ recipient mice and harvested the spleen two days later. Consistent with previous studies^26–29^, transferred ILC3s localized predominantly to the white pulp at the border of the splenic marginal zone (**Figure 4A-B**). To directly compare the capacity of ILC3s and cDC2s to induce Ag-dependent CD4^+^ T-cell activation and proliferation *in vivo*, LCMV-GP_61-80_-loaded mature ILC3s or cDC2s were transferred *i.v.* into MHCII^-/-^ recipient mice. The following day, CTV-labeled naive Nur77-GFP SMARTA CD4^+^ T cells were injected *i.v.*, and T-cell proliferation was assessed two days later (**Figure 4C**). Both the frequency and absolute number of donor SMARTA CD4^+^ T cells were significantly higher in the spleen of mice receiving mature cDC2s than in those receiving mature ILC3s (**Figure 4D**). Despite this lower overall SMARTA accumulation, TCR-engaged SMARTA CD4^+^ T cells underwent an almost similar number of cell divisions following priming by mature ILC3s or cDC2s (**Figure 4E**). The progressive decrease in Nur77-GFP MFI across successive cell divisions was also comparable between ILC3- and cDC2-primed SMARTA CD4^+^ T cells, consistent with comparable TCR signaling dynamics (**Figure 4F-G**). Accordingly, T cells primed by either APC type showed effector and activation phenotypes, as evidenced by the expression of CD69, CD25, and PD-1 (**Figure 4H-J**). Taken together, transferred mature ILC3s localize to the splenic white pulp and induce CD4^+^ T-cell responses *in vivo*.

**Figure 4:**
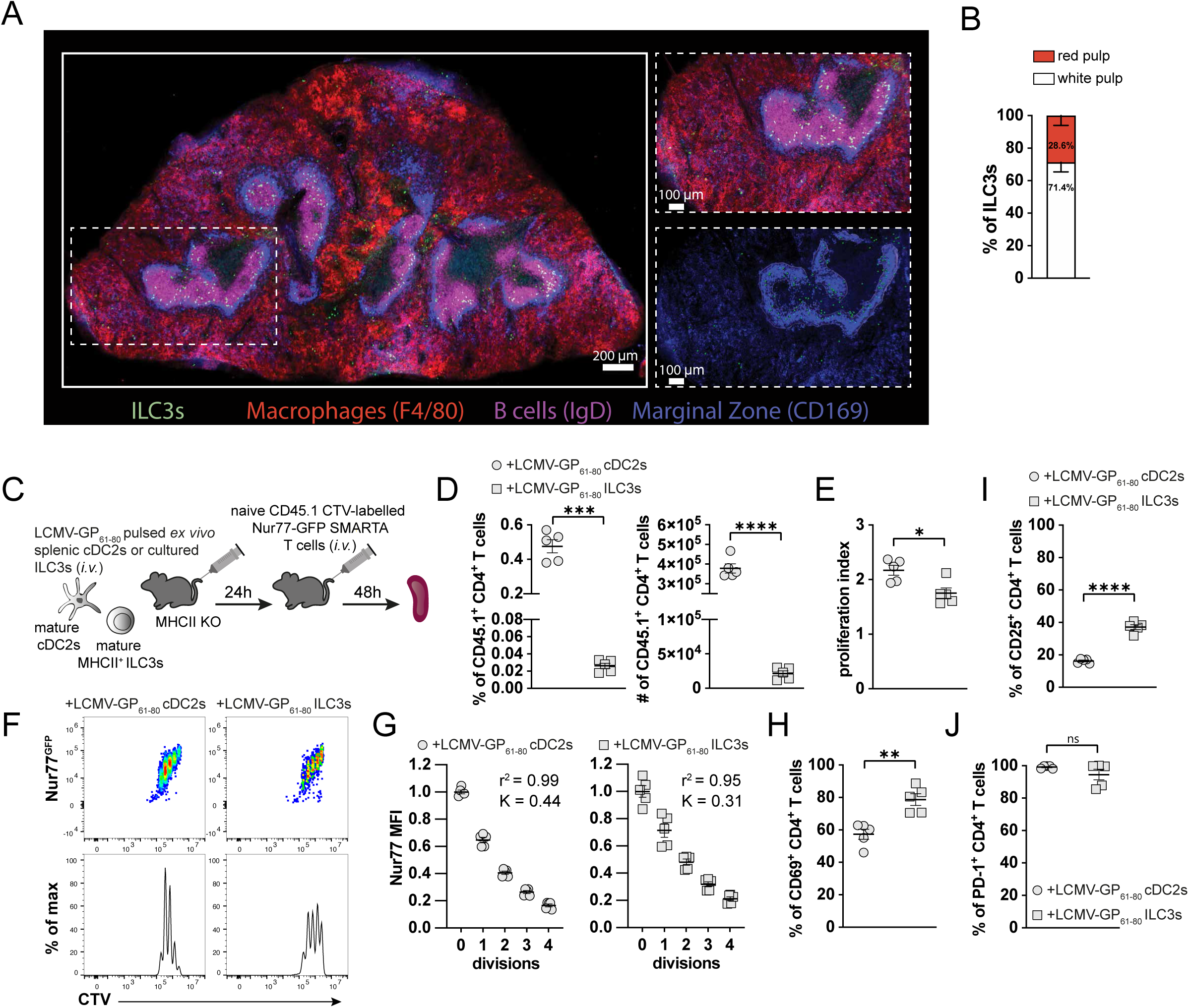
Mature ILC3s localize to the white pulp at the border of the splenic marginal zone and induce CD4^+^ T-cell proliferation. (**A**) Representative confocal image showing ILC3s (GFP; green), macrophages (F4/80; red), B cells (IgD; magenta), and the marginal zone (CD169; blue) in the spleen of MHCII^-/-^ recipient mice two days after transfer of mature SrtA^+^ ILC3s. Dashed box indicates the magnified region. Scale bars; 200 μm and 100 μm. (**B**) Quantification of the distribution of transferred ILC3s in the red and white pulp. Data are representative of three biological replicates. (**C**) Scheme of the experimental setup for the *in vivo* T-cell proliferation assay. (**D**) Frequencies and absolute numbers of Nur77-GFP SMARTA CD45.1^+^CD4^+^ T cells in the spleen. (**E**) Quantification of the proliferation index of Nur77-GFP SMARTA CD45.1^+^CD4^+^ T cells in the spleen. Representative FACS proliferation profiles (**F**) and Nur77-GFP MFI (**G**) across successive cell divisions of activated Nur77-GFP SMARTA CD45.1^+^CD4^+^ T cells in the spleen. Frequencies of CD69^+^ (**H**), CD25^+^ (**I**), and PD-1^+^ (**J**) Nur77-GFP SMARTA CD45.1^+^CD4^+^ T cells in the spleen. Mean values ± SEM of 5 mice from two independent experiments. Statistical significance was determined by unpaired Welch’s Test. p>0.05 (ns; not significant), p<0.05 (*), p <0.01 (**), p <0.001 (***) and p < 0.0001 (****).

### Tracing Ag-specific interaction between adoptively transferred APCs and T cells *in vivo*

To determine whether LIPSTIC could label naive CD4⁺ T cells following cognate interactions with ILC3s or cDC2s in the spleen, we first generated naive G5-hCD2⁺ SMARTA CD4⁺ T cells by reconstituting CD45.1/CD45.2 recipient mice with G5-hCD2⁺ NUPA10hd SMARTA progenitors (**Figure S4A**). Reconstituted mice exhibited normal frequencies of donor lymphoid progenitor populations, thymocyte subsets, spleen size and naive (CD62L^+^CD44^-^) T-cell populations in the spleen (**Figure S4B-H**). Next, mature SrtA-hCD58^+^ cDC2s or ILC3s were loaded with either the cognate LCMV-GP_61–80_ or with a control OVA_323–339_ peptide and injected *i.v.* into MHCII^-/-^ recipient mice. The following day, naive G5-hCD2^+^ SMARTA CD4^+^ T cells were injected *i.v.*, followed by three *i.p.* injections of biotin-LPETG (**Figure 5A**). Flow cytometric analysis of splenocytes of the MHCII^-/-^ recipient mice revealed LIPSTIC labeling of transferred SMARTA CD4^+^ T cells by LCMV-GP_61-80_-loaded cDC2s and ILC3s (**Figure 5B-I**). The frequency of biotin^+^CD69^+^ SMARTA CD4^+^ T cells was markedly reduced following interactions with ILC3s compared with cDC2s (**Figure 5E,I**). No LIPSTIC labeled SMARTA CD4^+^ T cells were detected in the inguinal LNs (iLNs), suggesting that the priming was confined to the spleen (**Figure 5B,F; right**). Together, these results demonstrate that the LIPSTIC hCD58/hCD2 receptor-ligand pair enables the detection of CD4^+^ T cells following Ag-specific interactions with both conventional cDC2s and unconventional ILC3s *in vivo*.

**Figure 5:**
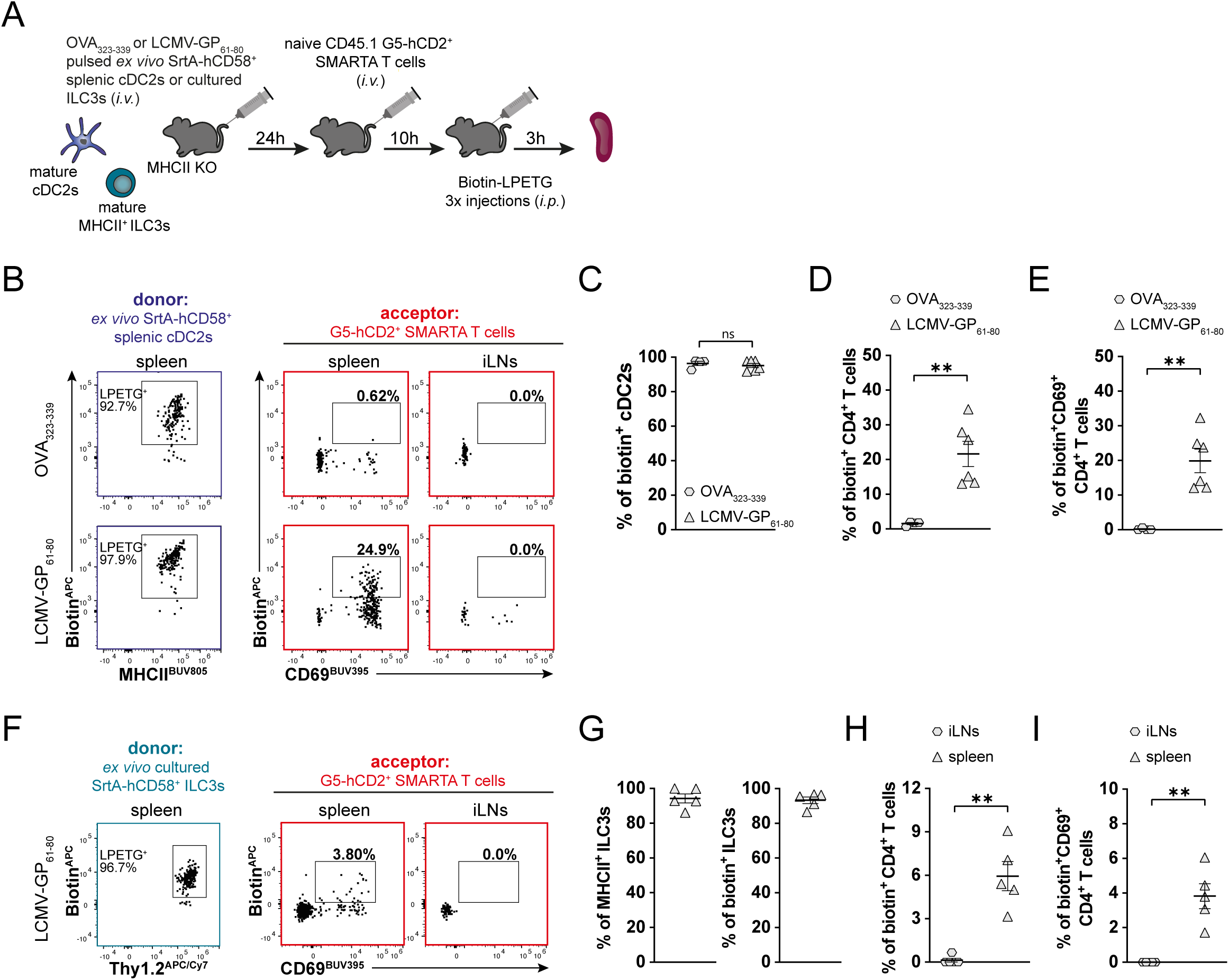
Tracing Ag-specific interaction of adoptively transferred ILC3s with T cells in the spleen. (**A**) Experimental setup for the LIPSTIC labeling of G5-hCD2^+^ T cells by mature SrtA-hCD58^+^ cDC2s or ILC3s in the spleen. (**B**) Representative flow cytometry dot plots depicting the loading of biotin-LPETG on mature SrtA-hCD58^+^ cDC2s (left) and its transfer to G5-hCD2^+^ T cells in the spleen (middle) and inguinal lymph nodes (iLNs; right) 13 h post T-cell transfer. Frequencies of biotin^+^ cDC2s (**C**), biotin^+^ (**D**) and biotin^+^CD69^+^ (**E**) CD4^+^ T cells in the spleen. Mean values ± SEM of 4-6 mice from two independent experiments. (**F**) Representative flow cytometry dot plots depicting the loading of biotin-LPETG on mature SrtA-hCD58^+^ ILC3s (left) and its transfer to G5-hCD2^+^ T cells in the spleen (middle) and iLNs (right) 13 h post T-cell transfer. (**G**) Frequencies of MHCII^+^ and biotin^+^ ILC3s in the spleen. Frequencies of biotin^+^ (**H**) and biotin^+^CD69^+^ (**I**) CD4^+^ T cells in the spleen and iLNs. Mean values ± SEM of 5 mice from three independent experiments. Statistical significance was determined by unpaired Welch’s Test. p>0.05 (ns; not significant) and p <0.01 (**).

## Discussion

In this study, we established and validated a versatile LIPSTIC system to selectively label and recover Ag-specific CD4⁺ T cells following interactions with distinct APC subsets *in vivo*. Central to this is the use of the human CD58/CD2 receptor-ligand pair fused to SrtA as donor and G5 tag as acceptor, respectively. Our implementation utilizes NUPA10hd^+^ immortalized hematopoietic progenitors to generate SrtA-hCD58⁺ DCs and G5-hCD2⁺ T cells, a strategy that circumvents the need for generating new genetically modified mouse lines and thus enables rapid application to various research models. We validated the system *in vitro* and *in vivo* by successfully detecting label transfer from DCs to CD4⁺ T cells in an Ag-dependent manner. We extended the use of this model to track T-cell interactions with unconventional APCs, such as ILC3s. This platform establishes a new framework for characterizing how various APC subsets, both professional and non-professional, drive Ag-specific T-cell responses.

The original LIPSTIC mouse model used SrtA-tagged CD40L expressed by T cells to label G5-CD40^+^ APCs. Although we have successfully achieved the reverse by swapping SrtA and G5 tags and observe LIPSTIC labeling in a cell line, retroviral overexpression of CD40/CD40L and subsequent adoptive transfer *in vivo* would lead to sustained, Ag-independent immune activation. While the development of knock-in mice expressing the reversed SrtA-CD40 and CD40L-G5 pair is technically feasible, such an approach would fundamentally restrict the system’s utility to cell populations that inherently express CD40 or CD40L. The integration of the NUPA10hd⁺ progenitor system with the immunologically inert human CD58/CD2 receptor-ligand pair provides a highly efficient method for studying cell-cell interaction in any direction allowing for rapid adaptation to different immune cell lineages.

While adoptively transferred ILC3s successfully induce Ag-specific activation and proliferation in CD4⁺ T cells, LIPSTIC labeling and the T-cell response differed from those observed with cDC2s. This disparity may be attributed to morphological differences, such as the dendritic processes of cDCs that expand the available surface area for T-cell encounters, as well as variations in cellular localization and co-stimulatory molecule expression. However, because *in vitro* labeling and *in vivo* Nur77 upregulation were comparable between both populations, the reduced labeling frequency does not necessarily imply an intrinsic deficiency in ILC3 Ag-presentation capacity. These findings potentially suggest that the physical frequency of interaction, rather than the qualitative efficiency of the encounter, may be the primary driver of the observed differences. Ultimately, further investigation is required to determine if these distinct APC subsets drive qualitatively different functional fates in Ag-specific T cells.

Beyond APC/T-cell interactions, the hCD58/hCD2 receptor-ligand LIPSTIC pair could be broadly applicable to studying diverse cell-cell interactions *in vivo*. For example, Lee *et al.* recently adapted a LIPSTIC strategy to distinguish proximal and distal daughter CAR T cells after their first division^30^, highlighting the potential of LIPSTIC to link specific cell-cell interactions to the subsequent fate and function of the interacting cells. Likewise, our system could be used to track the survival, phenotype, migration, and transcriptional state of G5-hCD2⁺ CAR T cells following interactions with SrtA-hCD58⁺ target cells. Moreover, combining differently labeled LPETG substrates could potentially enable temporal mapping of sequential interactions between individual cells and multiple APC or tumor cell populations. Despite its broad utility, several limitations of our approach should be considered. The persistence of LIPSTIC labeling on acceptor cells remains unknown, potentially limiting long-term tracing studies. In addition, the current approach relies on the adoptive transfer of genetically engineered cells, which may restrict its applicability to rare or tissue-resident cell populations. Nevertheless, the combination of the novel LIPSTIC receptor-ligand pair together with a robust *in vitro* differentiation toolkit provides a versatile and adaptable system for tracing cell-cell interactions within the immune system.

## Acknowledgments

We thank Martha Gaio for the critical reading of the manuscript. We would also like to thank Dr. Gaël Auray from the DBM FACS Facility (Basel) for the cell sorting. Finally, we thank the staff of the Animal Facility Mattenstrasse for the animal work. Cartoons (Figure 1) were created using Biorender.com. The remaining cartoons were adapted from our previous publication^20^ under a CC BY license, originally designed by Nadine Anslinger.

## Authors contribution

GPBS: Funding acquisition, Writing – original draft, Conceptualization, Methodology, Validation, Investigation, Visualization.

YF: Writing - Review & Editing, Conceptualization, Methodology, Validation, Investigation, Visualization. AL: Writing - Review & Editing, Conceptualization, Investigation.

EH: Writing - Review & Editing, Investigation.

DF: Funding acquisition, Writing - Review & Editing, Conceptualization.

GT: Funding acquisition, Writing - Review & Editing, Conceptualization, Methodology, Investigation.

