## Supplementary material for "Cognate T-cell labeling by conventional and unconventional antigen-presenting cells *in vivo* using a modular SrtA-based toolkit": Supplemenatry material

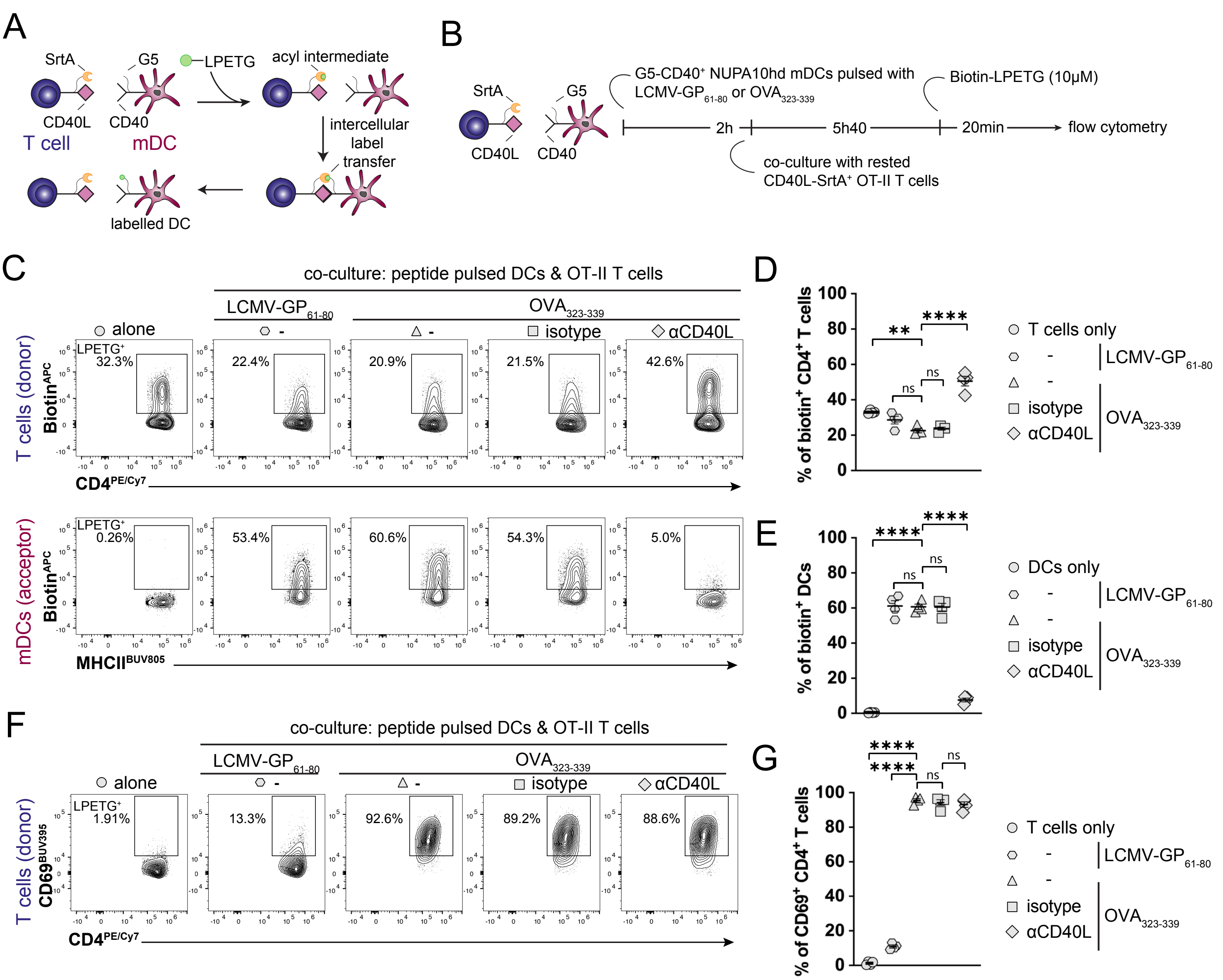
A
SrtA
G5
LPETG
acyl intermediate
CD40L
CD40
T cell
mDC
intercellular label transfer
labelled DC
B
SrtA
G5
CD40L
CD40
G5-CD40+ NUPA10hd mDCs pulsed with LCMV-GP61-80 or OVA323-339
Biotin-LPETG (10μM)
2h
5h40
20min
flow cytometry
co-culture with rested CD40L-SrtA+ OT-II T cells
C
co-culture: peptide pulsed DCs & OT-II T cells
LCMV-GP61-80
OVA323-339
alone
-
-
isotype
αCD40L
T cells (donor)
BiotinAPC
LPETG+
32.3%
22.4%
20.9%
21.5%
42.6%
CD4PE/Cy7
mDCs (acceptor)
BiotinAPC
LPETG+
0.26%
53.4%
60.6%
54.3%
5.0%
MHCIIBUV805
D
% of biotin+ CD4+ T cells
**
****
ns
ns
T cells only
-
-
isotype
αCD40L
LCMV-GP61-80
OVA323-339
E
% of biotin+ DCs
****
****
ns
ns
DCs only
-
-
isotype
αCD40L
LCMV-GP61-80
OVA323-339
F
co-culture: peptide pulsed DCs & OT-II T cells
LCMV-GP61-80
OVA323-339
alone
-
-
isotype
αCD40L
T cells (donor)
CD69BUV395
LPETG+
1.91%
13.3%
92.6%
89.2%
88.6%
CD4PE/Cy7
G
% of CD69+ CD4+ T cells
****
****
ns
ns
T cells only
-
-
isotype
αCD40L
LCMV-GP61-80
OVA323-339

**Figure S1: LIPSTIC labeling of G5-CD40$^+$ mDCs by CD40L-SrtA$^+$ T cells *in vitro*.** (**A**) Scheme of the LIPSTIC approach. (**B**) Experimental setup to analyze intercellular labeling of G5-CD40$^+$ NUPA10hd mature DCs (mDCs) by CD40L-SrtA$^+$ OT-II CD4$^+$ T cells pre-treated or not with a blocking αCD40L or isotype antibody. (**C**) Representative flow cytometry contour plots of biotin labeling of mDCs (intercellular transfer) and CD4$^+$ T cells (acyl intermediate). (**D**) Frequency of biotin$^+$ CD4$^+$ T cells. (**E**) Frequency of biotin$^+$ mDCs. (**F**) Representative flow cytometry contour plots of CD69 expression on CD4$^+$ T cells. (**G**) Frequency of CD69$^+$ CD4$^+$ T cells. Mean values ± SEM of four independent experiments. Statistical significance was determined by ordinary one-way ANOVA, followed by Tukey's multiple-comparisons test. $p>0.05$ (ns; not significant), $p < 0.01$ (**) and $p < 0.0001$ (****).

Figure S2

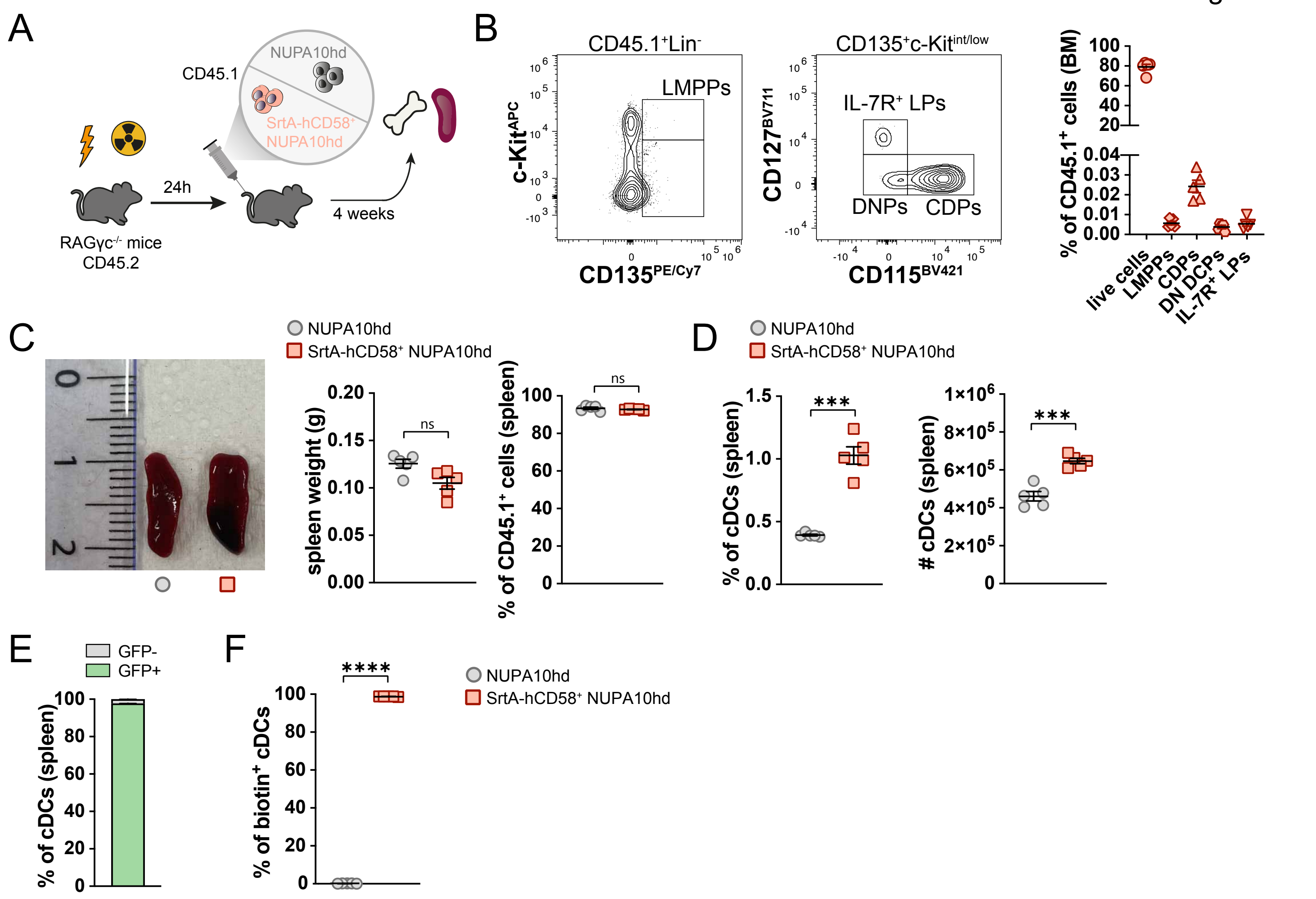

A
CD45.1
NUPA10hd
SrtA-hCD58+ NUPA10hd
24h
4 weeks
RAGγc-/- mice
CD45.2
B
CD45.1+Lin-
LMPPs
c-KitAPC
CD135PE/Cy7
CD135+c-Kitint/low
IL-7R+ LPs
DNPs
CDPs
CD127BV711
CD115BV421
% of CD45.1+ cells (BM)
live cells
LMPPs
CDPs
DN DCPs
IL-7R+ LPs
C
NUPA10hd
SrtA-hCD58+ NUPA10hd
spleen weight (g)
ns
% of CD45.1+ cells (spleen)
ns
D
NUPA10hd
SrtA-hCD58+ NUPA10hd
% of cDCs (spleen)
***
### cDCs (spleen)
***
E
GFP-
GFP+
% of cDCs (spleen)
F
****
% of biotin+ cDCs
NUPA10hd
SrtA-hCD58+ NUPA10hd

**Figure S2: SrtA-hCD58$^+$ NUPA10hd progenitors give rise to functional SrtA-hCD58$^+$ splenic cDCs in RAGγc$^{-/-}$ reconstituted mice.** (**A**) Scheme of the experimental setup for the *in vivo* reconstitution of Hox progenitors. (**B**) Gating strategy and frequencies of donor CD45.1$^+$ cells in bone marrow (BM) as well as CD45.1$^+$ lymphoid-primed multipotent progenitors (LMPPs), common DC progenitors (CDPs), double negative DC progenitors (DN DCPs) and IL-7R$^+$ lymphoid progenitors (LPs) at 4 weeks post-transfer in SrtA-hCD58$^+$ NUPA10hd mice. Lineage (Lin) cocktail consisted of antibodies against CD3, CD4, CD8, Ter119, CD11c, NK1.1, F4/80, B220, CD19 and Ly-6G. (**C**) Photography depicting spleens harvested from wild-type NUPA10hd and SrtA-hCD58$^+$ NUPA10hd mice (left). Quantification of the weight of the spleens (middle). Frequency of donor CD45.1$^+$ cells in the spleen (right). (**D**) Frequency and absolute number of CD45.1$^+$ cDCs (CD3$^-$CD19$^-$Ly6G$^-$F4/80$^-$CD11c$^+$MHCII$^+$) in the spleen. (**E**) Frequencies of GFP$^{+/-}$ CD45.1$^+$ cDCs in the spleen of SrtA-hCD58$^+$ NUPA10hd mice. (**F**) Frequency of CD45.1$^+$ biotin$^+$ cDCs among splenocytes from wild-type and SrtA-hCD58$^+$ NUPA10hd mice after *ex vivo* incubation with 100 µM biotin-LPETG for 30 min at 37°C. Mean values ± SEM of 5 mice from two independent experiments. Statistical significance was determined by unpaired Welch's Test. $p>0.05$ (ns; not significant), $p <0.001$ (***) and $p < 0.0001$ (****).

Figure S3

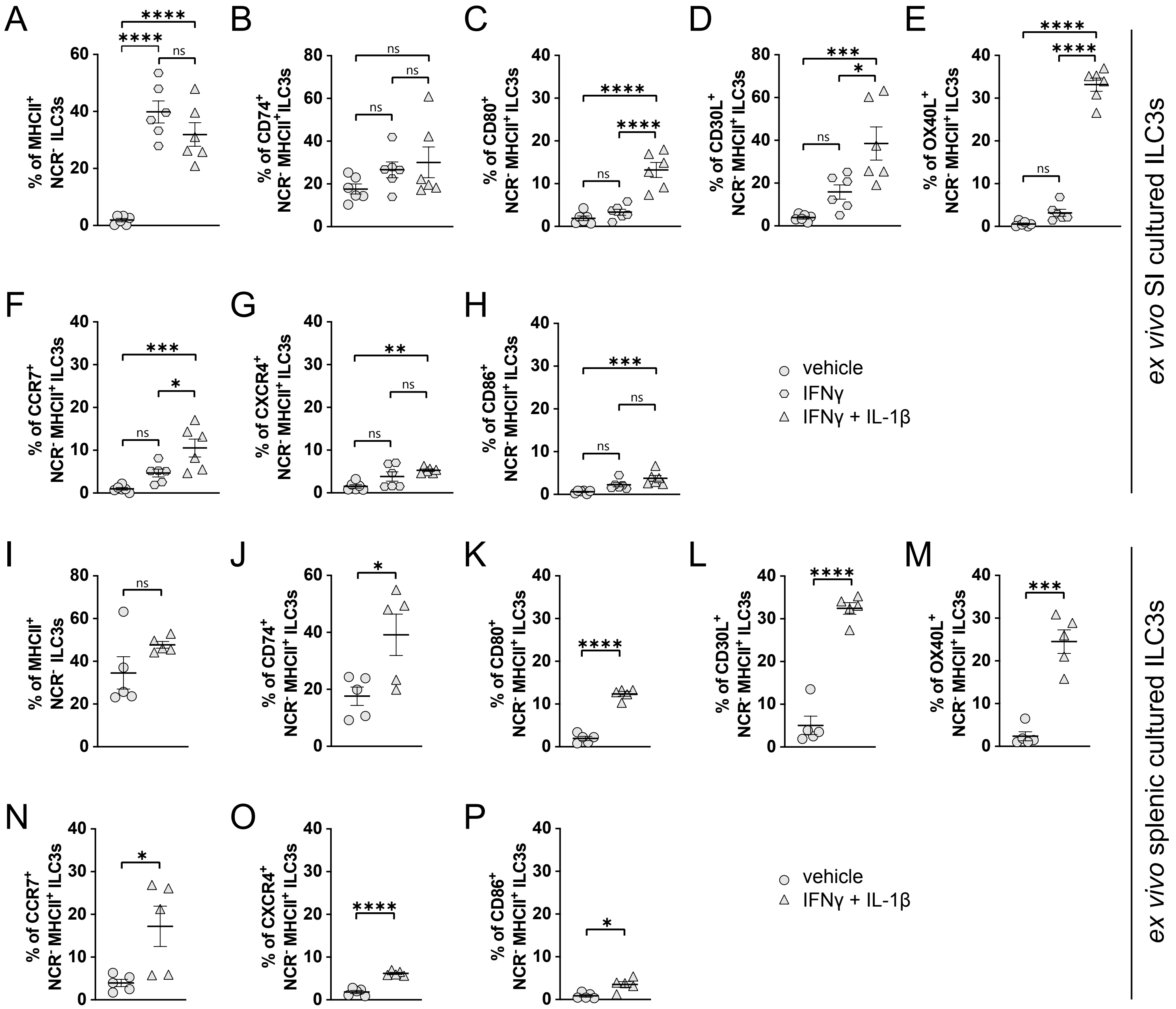
A
% of MHCII+ NCR- ILC3s
B
% of CD74+ NCR- MHCII+ ILC3s
C
% of CD80+ NCR- MHCII+ ILC3s
D
% of CD30L+ NCR- MHCII+ ILC3s
E
% of OX40L+ NCR- MHCII+ ILC3s
F
% of CCR7+ NCR- MHCII+ ILC3s
G
% of CXCR4+ NCR- MHCII+ ILC3s
H
% of CD86+ NCR- MHCII+ ILC3s
vehicle
IFNγ
IFNγ + IL-1β
ex vivo SI cultured ILC3s
I
% of MHCII+ NCR- ILC3s
J
% of CD74+ NCR- MHCII+ ILC3s
K
% of CD80+ NCR- MHCII+ ILC3s
L
% of CD30L+ NCR- MHCII+ ILC3s
M
% of OX40L+ NCR- MHCII+ ILC3s
N
% of CCR7+ NCR- MHCII+ ILC3s
O
% of CXCR4+ NCR- MHCII+ ILC3s
P
% of CD86+ NCR- MHCII+ ILC3s
vehicle
IFNγ + IL-1β
ex vivo splenic cultured ILC3s

**Figure S3: IL-1β induce a comparable maturation state in *ex vivo*-expanded splenic and small intestine (SI)-derived ILC3s.** Frequency of $MHCII^{+}NCR^{-}$ cells among *ex vivo*-expanded SrtA-hCD58$^{+}$ SI- (**A**) and splenic- (**I**) derived ILC3s stimulated with IFNγ only, IFNγ + IL-1β (SI) or IFNγ + IL-1β (splenic) or left unstimulated. $MHCII^{+}NCR^{-}$ ILC3s were defined as $GFP^{+}Thy1.2^{+}ROR\gamma t^{+}NKp46^{-}MHCII^{+}$ cells. Frequencies of CD74$^{+}$ (**B,J**), CD80$^{+}$ (**C,K**), CD30L$^{+}$ (**D,L**), OX40L$^{+}$ (**E,M**), CCR7$^{+}$ (**F,N**), CXCR4$^{+}$ (**G,O**), and CD86$^{+}$ (**H,P**) cells among SrtA-hCD58$^{+}$ $MHCII^{+}NCR^{-}$ SI- (**B-H**) and splenic- (**J-P**) derived ILC3s under the indicated conditions. Data are mean ± SEM from six (SI) and five (splenic) independent experiments. Statistical significance was assessed by one-way ANOVA with Tukey's multiple-comparisons test (SI) or unpaired Welch's Test (splenic). $p>0.05$ (ns; not significant), $p<0.05$ (*), $p < 0.01$ (**), $p <0.001$ (***) and $p < 0.0001$ (****).

Figure S4

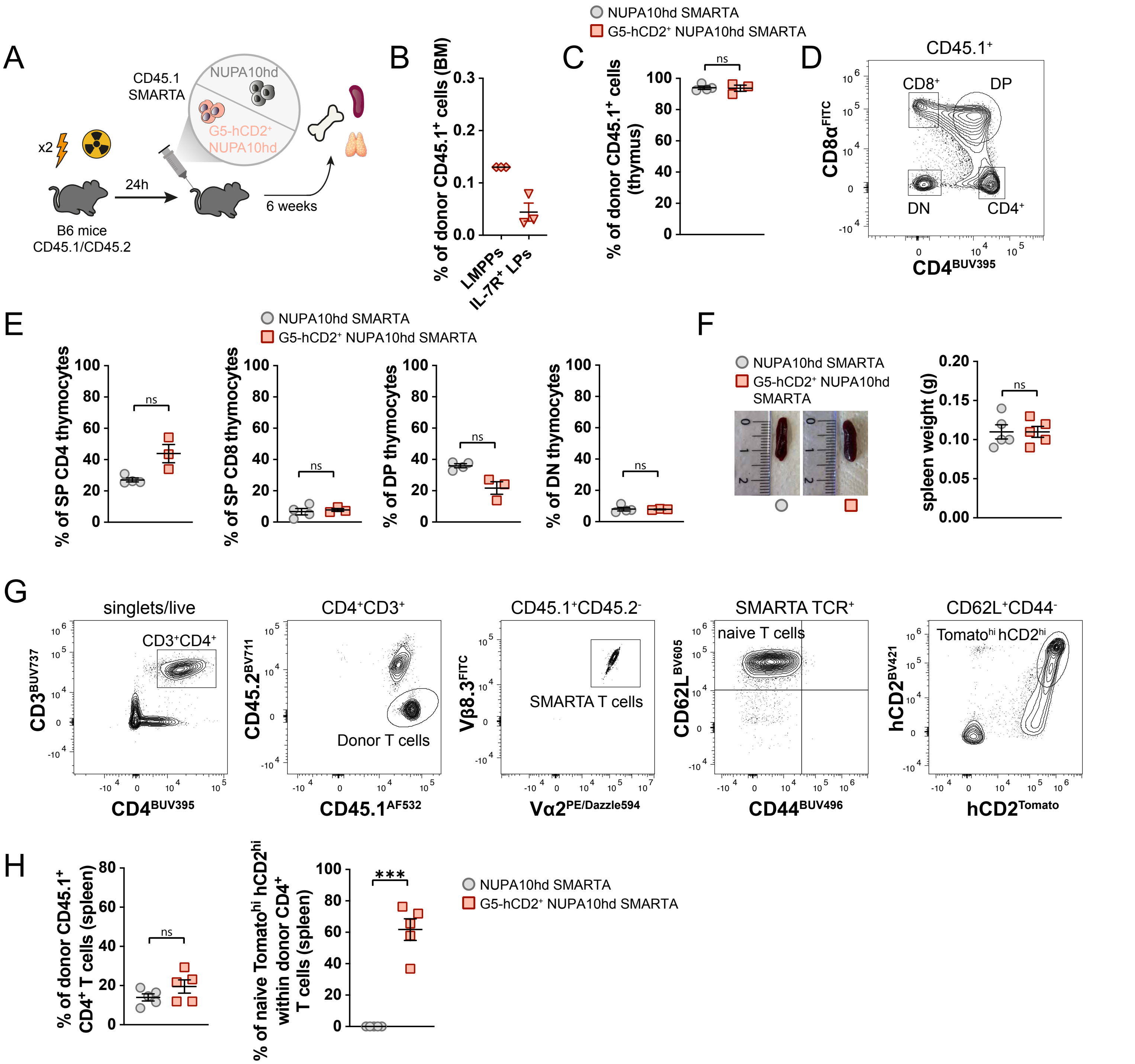
A
CD45.1
SMARTA
NUPA10hd
G5-hCD2+ NUPA10hd
x2
24h
6 weeks
B6 mice
CD45.1/CD45.2
B
% of donor CD45.1+ cells (BM)
LMPPs
IL-7R+ LPs
NUPA10hd SMARTA
G5-hCD2+ NUPA10hd SMARTA
C
% of donor CD45.1+ cells (thymus)
ns
D
CD45.1+
CD8+
DP
DN
CD4+
CD8α FITC
CD4 BUV395
E
% of SP CD4 thymocytes
% of SP CD8 thymocytes
% of DP thymocytes
% of DN thymocytes
F
spleen weight (g)
G
singlets/live
CD3+CD4+
CD3 BUV737
CD4 BUV395
CD4+CD3+
CD45.2 BV711
CD45.1 AF532
Donor T cells
CD45.1+CD45.2-
Vβ8.3 FITC
Vα2 PE/Dazzle594
SMARTA T cells
SMARTA TCR+
naive T cells
CD62L BV605
CD44 BUV496
CD62L+CD44-
Tomatohi hCD2hi
hCD2 BV421
hCD2 Tomato
H
% of donor CD45.1+ CD4+ T cells (spleen)
% of naive Tomatohi hCD2hi within donor CD4+ T cells (spleen)
***

**Figure S4: G5-hCD2$^+$ NUPA10hd SMARTA progenitors give rise to functional naive G5-hCD2$^+$ CD4$^+$ T cells in the spleen of CD45.1/CD45.2-reconstituted mice.** (**A**) Scheme of the experimental setup for the *in vivo* reconstitution of Hox progenitors. (**B**) Frequencies of donor CD45.1$^+$ lymphoid-primed multipotent progenitors (LMPPs) and IL-7R$^+$ lymphoid progenitors (LPs) in the bone marrow (BM) at 6 weeks post-transfer in G5-hCD2$^+$ NUPA10hd SMARTA mice. Lineage (Lin) cocktail consisted of antibodies against CD3, CD4, CD8, Ter119, CD11c, NK1.1, F4/80, B220, CD19 and Ly-6G. (**C**) Frequencies of donor CD45.1$^+$ cells in the thymus. (**D**) Gating strategy to determine CD45.1$^+$ single positive (SP) CD4$^+$, SP CD8$^+$, double positive (DP) and double negative (DN) thymocytes. (**E**) Frequencies of CD45.1$^+$ SP CD4$^+$, SP CD8$^+$, DP and DN thymocytes at 6 weeks post-transfer in wild-type and G5-hCD2$^+$ NUPA10hd SMARTA mice. (**F**) Photography depicting spleens harvested from wild-type and G5-hCD2$^+$ NUPA10hd SMARTA mice at 6 weeks post-transfer as well as the quantification of the weight of the spleens. (**G**) Gating strategy to determine naive donor CD45.1$^+$ Tomato$^{hi}$ hCD2$^{hi}$ CD4$^+$ T cells in the spleen. (**H**) Frequencies of donor CD45.1$^+$ CD4$^+$ T cells and of naive Tomato$^{hi}$ hCD2$^{hi}$ within donor CD4$^+$ T cells in the spleen at 6 weeks post-transfer in wild-type and G5-hCD2$^+$ NUPA10hd SMARTA mice. Mean values ± SEM of 3-5 mice from at least two independent experiments. Statistical significance was determined by unpaired Welch's Test. $p>0.05$ (ns; not significant) and $p<0.001$ (***).

**Table 1 : List of antibodies**

| Antibodies | Company | Catalog# |
|---|---|---|
| APC-conjugated anti-mouse CCR7 (4B12) | Thermo Fisher Scientific | Cat#17-1971-82 |
| APC-conjugated anti-biotin (REA746) | Mitenyi Biotec | Cat#130-110-952 |
| APC-conjugated Streptavidin | BioLegend | Cat#405207 |
| BV421-conjugated Streptavidin | BioLegend | Cat#405226 |
| PE/Cy7-conjugated anti-mouse CD4 (RM4-5) | BioLegend | Cat#100528 |
| BUV395-conjugated anti-mouse CD69 (HI.2F3) | Waters Biosciences | Cat#740220 |
| BUV805-conjugated anti-mouse I-A/I-E (M5/114.15.2) | Waters Biosciences | Cat#748844 |
| BV421-conjugated anti-mouse CD11c (N418) | BioLegend | Cat#117329 |
| APC/Fire750-conjugated anti-mouse CD69 (HI.2F3) | BioLegend | Cat#104548 |
| BV605-conjugated anti-mouse CD11c (N418) | BioLegend | Cat#117333 |
| APC-conjugated anti-mouse I-A/I-E (M5/114.15.2) | BioLegend | Cat#107613 |
| BUV737-conjugated anti-mouse CD3ε (17A2) | Waters Biosciences | Cat#612803 |
| BV421-conjugated anti-mouse I-A/I-E (M5/114.15.2) | BioLegend | Cat#107631 |
| APC/Cy7-conjugated anti-mouse CD90.2 (Thy1.2; 30-H12) | BioLegend | Cat#105327 |
| BUV395-conjugated anti-mouse CD4 (GK1.5) | Waters Biosciences | Cat#565974 |
| BUV496-conjugated anti-mouse CD44 (IM7) | Waters Biosciences | Cat#569706 |
| BV605-conjugated anti-mouse CD62L (MEL-14) | BioLegend | Cat#104437 |
| PerCP/Cy5.5-conjugated anti-mouse CD62L (MEL-14) | BioLegend | Cat#104432 |
| BV785-conjugated anti-mouse CD279 (PD-1; 29F.1A12) | BioLegend | Cat#135225 |
| AF532-conjugated anti-mouse CD45.1 (A20) | Thermo Fisher Scientific | Cat#58-0453-82 |
| PE-conjugated anti-mouse CD25 (PC61) | BioLegend | Cat#102007 |
| BUV395-conjugated anti-mouse CD11b (M1/70) | Waters Biosciences | Cat#563553 |
| BUV805-conjugated anti-mouse CD48 (HM48-1) | Waters Biosciences | Cat#741945 |
| BV421-conjugated anti-mouse CD115 (AFS98) | BioLegend | Cat#135513 |
| BV711-conjugated anti-mouse CD127 (IL-7Rα; A7R34) | BioLegend | Cat#135035 |
| BV786-conjugated anti-mouse Sca-1 (D7) | BioLegend | Cat#108139 |
| PE/Dazzle 594-conjugated anti-mouse CD150 (TC15-12F12.2) | BioLegend | Cat#115935 |
| PE/Cy7-conjugated anti-mouse CD135 (Flt-3;A2F10) | Waters Biosciences | Cat#567594 |
| APC-conjugated anti-mouse CD117 (cKit;2B8) | BioLegend | Cat#105812 |
| APC/Cy7-conjugated anti-mouse Ly6C (HK1.4) | BioLegend | Cat#128026 |
| PE-conjugated anti-mouse CD3ε (17A2) | BioLegend | Cat#100205 |
| PE-conjugated anti-mouse CD8α (53-6.7) | BioLegend | Cat#100707 |
| PE-conjugated anti-mouse Ter119 (Ter119) | BioLegend | Cat#116207 |
| PE-conjugated anti-mouse Ly6G (1A8) | BioLegend | Cat#127607 |
| PE-conjugated anti-mouse NK1.1 (PK136) | BioLegend | Cat#156503 |
| PE-conjugated anti-mouse CD11c (N418) | BioLegend | Cat#117307 |
| PE-conjugated anti-mouse CD11b (M1/70) | BioLegend | Cat#101208 |
| PE-conjugated anti-mouse CD19 (6D5) | BioLegend | Cat#115507 |
| PE-conjugated anti-mouse F4/80 (BM8) | BioLegend | Cat#111603 |

| | | |
|---|---|---|
| PE-conjugated anti-mouse B220 (RA3-6B2) | BioLegend | Cat#103207 |
| BUV496-conjugated anti-mouse I-A/I-E (M5/114.15.2) | Waters Biosciences | Cat#750281 |
| BUV563-conjugated anti-mouse CD11c (N418) | Waters Biosciences | Cat#749040 |
| BUV805-conjugated anti-mouse Ly6G (1A8) | Waters Biosciences | Cat#741994 |
| BV421-conjugated anti-mouse XCR1 (ZET) | BioLegend | Cat#148216 |
| PerCP/Cy5.5-conjugated anti-mouse CD172 (P84) | BioLegend | Cat#144009 |
| PE/Cy7-conjugated anti-mouse Ly6C (HK1.4) | BioLegend | Cat#128018 |
| APC/Cy7-conjugated anti-mouse F4/80 (BM8) | BioLegend | Cat#123118 |
| PE-conjugated anti-mouse TCRγδ (GL3) | BioLegend | Cat#118107 |
| BUV805-conjugated anti-mouse KLRG1 (2F1) | Waters Biosciences | Cat#741993 |
| BV421-conjugated anti-mouse CCR6 (140706) | Waters Biosciences | Cat#564736 |
| BV605-conjugated anti-mouse NK1.1 (PK136) | BioLegend | Cat#108739 |
| BV650-conjugated anti-mouse CD335 (NKp46; 29A1.4) | BioLegend | Cat#137635 |
| BV786-conjugated anti-mouse RORγt (Q31-378) | Waters Biosciences | Cat#564723 |
| BB700-conjugated anti-mouse GATA3 (L50-823) | Waters Biosciences | Cat#566643 |
| PE/Cy7-conjugated anti-mouse T-bet (4B10) | BioLegend | Cat#644823 |
| BUV395-conjugated anti-mouse CD74 (In-1) | Waters Biosciences | Cat#740274 |
| BV605-conjugated anti-mouse CD80 (16-10A1) | BioLegend | Cat#104729 |
| RB670-conjugated anti-mouse CXCR4 (2B11/CXCR4) | Waters Biosciences | Cat#771128 |
| RB744-conjugated anti-mouse CD30L (RM153) | Waters Biosciences | Cat#757484 |
| PE/Dazzle 594-conjugated anti-mouse CD86 (GL-1) | BioLegend | Cat#105041 |
| PE/Cy7-conjugated anti-mouse CD252 (OX40L; RM134L) | BioLegend | Cat#108813 |
| BUV496-conjugated anti-mouse CD45.2 (104) | Waters Biosciences | Cat#569670 |
| FITC-conjugated anti-mouse CD3ε (145-2C11) | BioLegend | Cat#100204 |
| FITC-conjugated anti-mouse CD8α (53-6.7) | Thermo Fisher Scientific | Cat#11-0081-82 |
| FITC-conjugated anti-mouse Ter119 (Ter119) | BioLegend | Cat#116215 |
| FITC-conjugated anti-mouse Ly6G (1A8) | BioLegend | Cat#127606 |
| FITC-conjugated anti-mouse NK1.1 (PK136) | Thermo Fisher Scientific | Cat#11-5941-85 |
| FITC-conjugated anti-mouse CD11c (N418) | BioLegend | Cat#117306 |
| FITC-conjugated anti-mouse F4/80 (BM8) | BioLegend | Cat#123107 |
| FITC-conjugated anti-mouse CD4 (GK1.5) | BioLegend | Cat#100406 |
| FITC-conjugated anti-mouse B220 (RA3-6B2) | BioLegend | Cat#103205 |
| PerCP/Cy5.5-conjugated anti-mouse CD45.1 (A20) | BioLegend | Cat#110725 |
| FITC-conjugated anti-mouse Vβ8.3 (1B3.3) | BioLegend | Cat#156305 |
| BV421-conjugated anti-human CD2 (RPA-2.10) | BioLegend | Cat#300229 |
| BV711-conjugated anti-mouse CD45.2 (A20) | BioLegend | Cat#109847 |
| PE/Dazzle 594-conjugated anti-mouse Vα2 (B20.1) | BioLegend | Cat#127828 |
| APC-conjugated anti-mouse CD8α (53-6.7) | BioLegend | Cat#100711 |
